# Reprogrammed mRNA translation drives resistance to therapeutic targeting of ribosome biogenesis

**DOI:** 10.1101/847723

**Authors:** E. P. Kusnadi, A. S. Trigos, C. Cullinane, D. L. Goode, O. Larsson, J. R. Devlin, K. T. Chan, D. P. De Souza, M. J. McConville, G. A. McArthur, G. Thomas, E. Sanij, G. Poortinga, R. D. Hannan, K. M. Hannan, J. Kang, R. B. Pearson

**Author notes:** These authors contributed equally (K. M. Hannan, J. Kang, R. B. Pearson). Corresponding Author: Richard B. Pearson, Peter MacCallum Cancer Centre, 305 Grattan St, Melbourne, Victoria, Australia, 3000.

## Abstract

Elevated ribosome biogenesis in oncogene-driven cancers is commonly targeted by DNA-damaging cytotoxic drugs. Our first-in-human trial of CX-5461, a novel, less genotoxic agent that specifically inhibits ribosome biogenesis via suppression of RNA Polymerase I (Pol I) transcription, revealed single agent efficacy in refractory blood cancers. Despite this clinical response, patients were not cured. In parallel, we demonstrated a marked improvement in the *in vivo* efficacy of CX-5461 in combination with PI3K/AKT/mTORC1 pathway inhibitors. Here we show that this improved efficacy is associated with specific suppression of translation of mRNAs encoding regulators of cellular metabolism. Importantly, acquired resistance to this co-treatment is driven by translational re-wiring that results in dysregulated cellular metabolism and induction of a cAMP-dependent pathway critical for the survival of blood cancers including lymphoma and acute myeloid leukemia. Our studies identify the molecular mechanisms underpinning the response of blood cancers to selective ribosome biogenesis inhibitors and identify metabolic vulnerabilities that will facilitate the rational design of more effective regimens for Pol I-directed therapies.

## Introduction

Many of the commonly used DNA-damaging cytotoxic drugs including 5-fluorouracil (5-FU), etoposide and oxaliplatin also interfere with ribosome biogenesis and target the “addiction” of transformed cells to elevated rates of ribosome biogenesis, inducing the “impaired ribosome biogenesis checkpoint” (IRBC) (*1*). Consequently, a number of less-genotoxic drugs that selectively target ribosome biogenesis and subsequently induce both the p53-dependent IRBC and/or p53-independent nucleolar-associated activation of the DNA-damage response have been developed and are showing increasing promise in clinical investigations (*1, 2*). We developed the “first in class” selective inhibitor of ribosome biogenesis, CX-5461, which targets RNA Polymerase I (Pol I) transcription, suppressing ribosomal RNA (rRNA) synthesis (*3*). Moreover, we demonstrated its *in vivo* efficacy in mouse models of both blood and prostate cancer (*3–7*). Critically, we recently completed a Phase I clinical trial of CX-5461, demonstrating single-agent efficacy in patients with advanced hematological malignancies (*8*). This dose escalation clinical study of 17 patients demonstrated that CX-5461 induced rapid on-target inhibition of Pol I transcription, resulting in prolonged partial response in one patient with anaplastic large cell lymphoma and stable disease in 5 patients with diffuse large B-cell lymphoma and myeloma (*8*).

In parallel studies, we reasoned that simultaneous targeting of ribosome biogenesis and protein synthesis, the critical processes where oncogenic networks converge to maintain cancer cell growth and survival, would improve the clinical benefit of CX-5461 and overcome many of the upstream potential drivers of resistance (*9–12*). Indeed, we demonstrated that combining CX-5461 with an inhibitor of PI3K/AKT/mTORC1-dependent mRNA translation, everolimus, markedly improved the pre-clinical therapeutic efficacy of either drug alone *in vivo* (*4, 6*). However, the mechanisms underlying this synergistic effect and the development of ensuing resistance to both CX-5461 alone or in combination with everolimus are unclear and their definition will be critical in optimizing the clinical efficacy of Pol I-directed ‘ribosome-targeting’ therapies.

To interrogate the molecular basis of the response to CX-5461 and the CX-5461 plus everolimus combination, we first performed genome-wide translational profiling to characterise the acute changes in mRNA usage of MYC-driven B-cell lymphomas in mice treated with CX-5461 and the mTORC1 inhibitor, everolimus. CX-5461 had minimal effect on the lymphoma’s translatome after treatment for 2 hours, however the combination therapy specifically inhibited the translation of mRNAs encoding multiple components of the translational apparatus and enzymes that regulate energy metabolism. The importance of this selective targeting of translation was emphasised by our finding that acquired resistance to the combination therapy was driven by upregulation of the translation of mRNAs encoding key components of the mitochondrial respiration network and the cAMP-EPAC1/2-RAP1 survival pathway. We confirmed the functional importance of this re-programming by demonstrating that both CX-5461 and combination therapy-resistant cells are more metabolically active and have elevated levels of cAMP. Importantly, specific inhibition of EPAC1/2 re-sensitizes resistant cells to the combinatorial ribosome-targeting therapy. More broadly, EPAC1/2 are also elevated in human acute myeloid leukemia (AML) and inhibition of EPAC1/2 reduced the viability of AML cell lines, demonstrating the critical role of this pro-survival pathway in lymphoma and AML. These studies reveal a key mechanism by which alterations in the translation of mRNAs encoding metabolic enzymes drive the response of tumor cells to ribosome-targeting therapies. They identify metabolic vulnerabilities that can be targeted to improve the response of hematological tumors to selective ribosome biogenesis inhibitors, which we predict will also be highly relevant for high proportion of cancers that are characterised by elevated ribosome biogenesis including cancers treated with the numerous standard of care cytotoxic drugs that inhibit this process.

## Results

### Acute inhibition of ribosome biogenesis and function selectively reduces translation of mRNAs encoding components of the translational apparatus

To examine the mechanisms of response to targeting ribosome biogenesis and protein synthesis by CX-5461 and everolimus, Eμ-*Myc* B-cell lymphoma (MSCV *Gfp*; clone #4242) cells were transplanted into C57BL/6 mice (*3, 4*). Lymphoma-bearing mice were treated on Day 10 post-transplant for 2 hours with CX-5461 (35 mg/kg), everolimus (EV; 5 mg/kg), or both (CX-5461+EV; 35 mg/kg CX-5461 and 5 mg/kg EV). We chose this early time point to exclude any confounding effects of drug-induced apoptosis on our molecular analyses. Indeed, no change in the percentage of GFP-positive lymphoma cells was observed in lymph nodes isolated from treated animals (Figure S1A). The on-target activity of EV was confirmed by the reduction in RPS6 phosphorylation (P-RPS6) (Ser240/244) in lymph nodes treated with EV or CX-5461+EV (Figure 1A, Figure S1B). In addition, despite the intrinsic variability in the p53 levels in the lymph nodes isolated from different mice, a significant increase of p53 protein was observed in response to CX-5461 and CX-5461+EV treatment as expected (Figure 1A, Figure S1C).

**Fig. 1:**
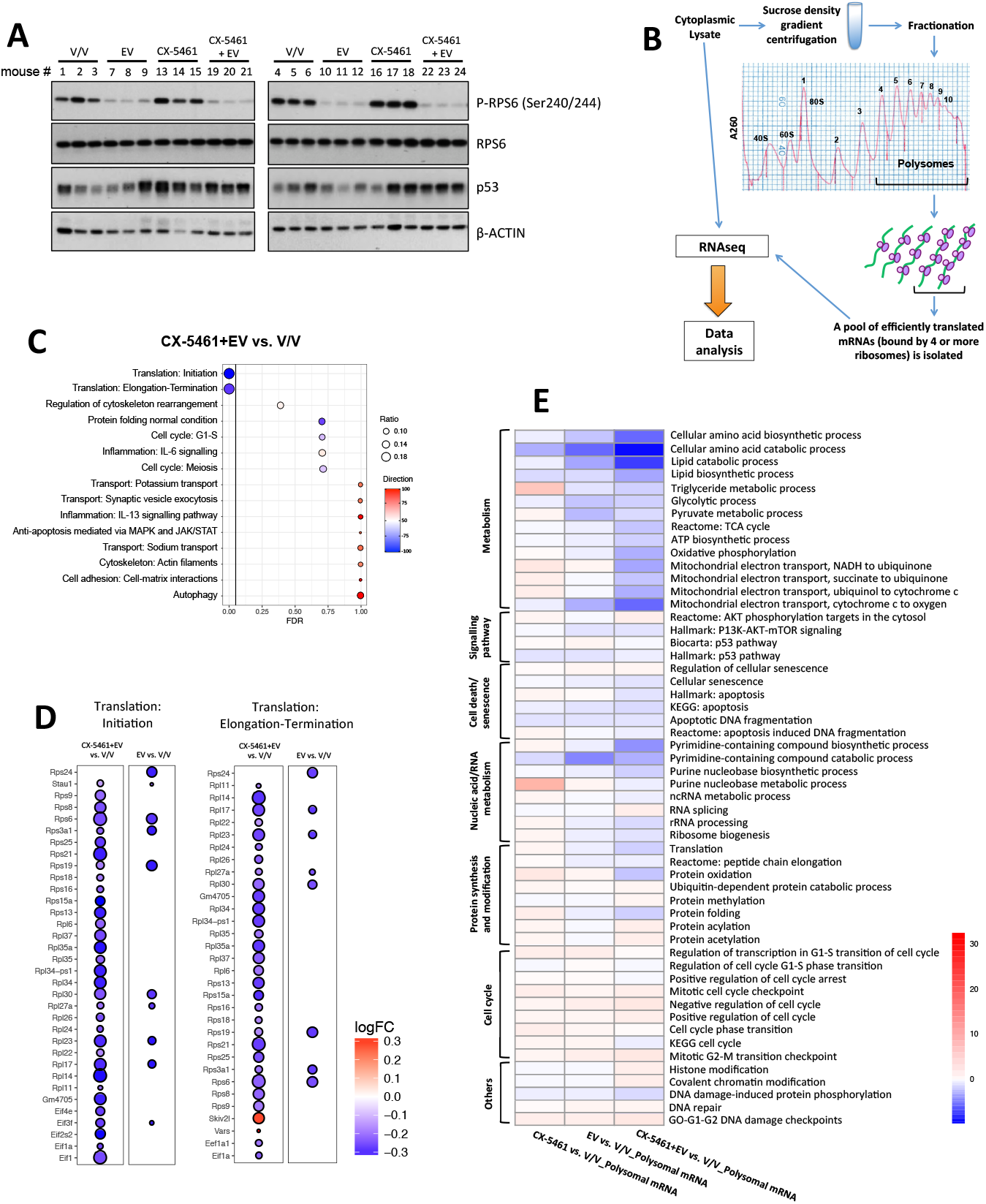
Acute inhibition of ribosome synthesis and function reduces translation of components of the translational machinery and decreases the abundance of polysome-associated, energy metabolism-related mRNAs. **(A)** Western analysis for on target effects for Everolimus (EV, P-RPS6) and CX-5461 (p53). Each lane represents equal amounts of protein from a single mouse that received drug vehicles (everolimus vehicle: 1% methylcellulose; CX-5461 vehicle: 25 mM NaH_2_PO_4_) (V/V; mouse #1-6), 5 mg/kg everolimus (EV; mouse #7-12), 35 mg/kg CX-5461 (mouse #13-18) or both drugs (CX-5461+EV; mouse #19-24) for 2 hours (n=6 per treatment group). Actin was used as a loading control. **(B)** Schematic of the polysome profiling analysis: cytoplasmic lysate was layered on top of a linear 10-40% sucrose gradient, ultracentrifuged (36,000 rpm, 2¼ hours, 4^0^C) and fractionated using the Foxy Jr Fraction Collector with constant monitoring of absorbance at 260 (A260) nm by an ISCO UA-6 Absorbance Detector. Fractions (one fraction per minute: 800 μL per tube) corresponding to polysomal mRNAs that were bound by four or more ribosomes were pooled and analyzed by RNA-seq followed by data analysis using anota2seq or limma. **(C)** Enrichment analysis by MetaCore® GeneGO of genes in “translation up” (red) and “translation down” (blue) categories identified by anota2seq analysis comparing lymph node cells isolated from mice in CX-5461+EV treatment group with the V/V group (n=6). “Ratio” is the value obtained by dividing the number of genes in our data assigned to a molecular process with the number of genes curated in the same process in MetaCore®’s database. logFC: log fold change; FDR: false discovery rate (adjusted *P* value). **(D)** Genes implicated in “*Translation: initiation*” and “*Translation: Elongation-Termination*” processes based on Figure 1C and Figure S1D. **(E)** Activity levels of key biological processes involved in cellular growth, proliferation and metabolism based on single sample gene set enrichment analysis (ssGSEA) of indicated comparisons. “Direction” visualises the percentage of genes associated with indicated pathways that are up- (red) or down- (blue) regulated. Data were obtained from n=6 mice per treatment group.

Given the specific alterations in translation patterns observed in response to mTOR inhibition (*13, 14*) and in genetic models of compromised ribosome biogenesis (*15*), we performed polyribosome (polysome) profiling to enable whole translatome analysis of mRNAs whose translation efficiency was altered by the acute ribosome-targeted combination therapy (Figure 1B). This transcriptome-wide translatomics approach involves the sequencing of polysome-associated mRNAs separated by sucrose density gradient ultracentrifugation accompanied by cytosolic mRNA sequencing (Figure 1B). To define changes in actively-translated mRNAs (indicated by attachment of four or more ribosomes) we compared the translational signatures of lymphoma cells isolated from combination therapy-treated mice (CX-5461+EV) versus vehicle-treated mice (V/V) using anota2seq analysis (*16*). After normalization to steady-state total RNA, the polysomal RNA-seq data indicated a robust reduction in the translation of transcripts encoding proteins involved in the regulation and activity of mRNA translation (Figure 1C). To further evaluate the significance of this translational re-wiring in mediating the synergy between CX-5461 and EV, we investigated the effects of CX-5461 and EV treatment as single agents. Not surprisingly, given the role of mTORC1 in translational control (*14, 17*), EV treatment alone affected the translation of mRNAs involved in mRNA translation (Figure S1D). In contrast, no significant effect on translation was observed upon single agent CX-5461 treatment (Figure S1E). However, quantitative comparison of the steady-state and polysome-associated mRNA counts in the top 2 enriched processes in Figure 1C and Figure S1D (“Translation initiation” and “Translation elongation”) revealed that the increased efficacy of combining EV with CX-5461 compared with EV alone was associated with specific and a more potent suppression of the translation of mRNAs encoding key proteins regulating mRNA translation (Figure 1D).

### Acute reduction in translation of the components of the translational apparatus is associated with decreased translation of energy metabolism-related mRNAs

Critically, despite this potent reduction in the efficiency of translation of components of the translational apparatus, cells treated with CX-5461+EV did not indiscriminately reduce the translation of all mRNAs globally. Single-sample gene set enrichment analysis (ssGSEA) (*18*) comparing abundance of all mRNAs associated with actively translating polysomes, rather than normalization to steady-state cytoplasmic mRNA abundance as previously performed in the anota2seq analysis, revealed a selective reduction in the translation of mRNAs encoding key metabolic processes in response to CX-5461+EV treatment. These mRNAs included those involved in energy metabolism, particularly in glycolysis and multiple steps of mitochondrial oxidative phosphorylation (oxphos) (Figure 1E). Moreover, analysis of the 5’UTRs of the targeted metabolic mRNAs revealed that they were 2 times more likely to be long (>200 nt; 17.4% vs. 8.5%) (Figure S1F), consistent with the reduced translation of translation initiation factors following CX-5461+EV treatment (Figure 1D), which are required for efficient translation of mRNAs with long 5’UTRs (*19*).

We hypothesised that this CX-5461+EV-induced targeting of metabolism was a key driver of the improved response to the combination. To test this hypothesis, we used metformin, a well-tolerated anti-diabetic drug that lowers cellular energy levels (*20*). Metformin treatment robustly increased cell death induced by CX-5461, consistent with a critical role for inhibition of metabolism in the improved efficacy of CX-5461+EV (Figure S1G). Moreover, metformin also markedly improved the therapeutic potency of CX-5461+EV (Figure S1H), emphasizing the importance of the targetable metabolic vulnerability in response to Pol I-directed therapy. Thus, we propose that the reduced translational activity in CX-5461+EV combination therapy-treated cells (Figure 1C), which selectively impaired translation of mRNAs encoding metabolic regulators, is a key mechanism in the synergistic effect of the two drugs and highlights the intimate coupling of mRNA translation and energy metabolism (*13, 21*).

We have shown that CX-5461 is efficacious in highly aggressive AML in both p53 WT and p53 null leukemic mice (*5*) that are also highly dependent on oxidative phosphorylation for survival (*22*). In order to establish whether energy metabolism is a potential metabolic vulnerability in other hematological cancers that could be exploited to enhance the efficacy of CX-5461, we evaluated the effects of metformin-CX-5461 combination in four human AML cell lines, representing a range of common oncogenic drivers of AML: MV4-11 (MLL-AF4 gene fusion), SHI-1 (MLL-AF6), SKM-1 (EZH2) and THP-1 (t(9;11) (*5*). As expected, metformin increased the abundance of phosphorylated AMPK and modestly reduced P-RPS6 levels in these cell lines, and everolimus promoted a robust decrease in the levels of P-RPS6 but had no effect on AMPK activation (Figure S1I). In MV4-11, SHI1 and THP-1 cells, a significant improvement of efficacy was observed when they were treated with CX-5461+metformin combination in comparison to the single agent treatment, (Figure S1J-M). Together, these data demonstrated the therapeutic potential of concurrent inhibition of energy metabolism and ribosome synthesis and function in haematological malignancies.

### Elevated energy metabolism is associated with resistance to ribosome targeting therapy

Despite the dramatic initial impact of the combination treatment on tumor growth, animals eventually succumb to lymphomagenesis (*3, 4*). Given that translation-dependent targeting of cellular metabolism is a key mediator of the acute response to combined targeting of ribosome biogenesis and mRNA translation (Figure 1C, 1D and 1E), we hypothesized that metabolic re-wiring driven by specific changes in mRNA translation would confer this resistance to therapy. To test this hypothesis, we established early-passage Eμ-*Myc* B-lymphoma cells lines from mice that were drug-naïve (from here onwards referred to as “CTRL” cells), previously treated with everolimus (“EV” cells), CX-5461 (“CX” cells) or CX-5461+EV (“CMB” cells) (Figure 2A). These early-passage cell lines were derived from lymph node extracts isolated from 12 different mice that were transplanted with Eμ-*Myc* B-lymphoma cells (clone #107) as indicated in Figure 2A. To confirm that CX and CMB cells maintained their drug resistance phenotypes, they were treated with EV, CX-5461 or CX-5461+EV *in vitro*. The drug-naive CTRL cell lines retained sensitivity to all treatments, while CX cells were resistant to CX-5461 and sensitive to CX-5461+EV and the CMB cells were unresponsive to all the treatments (Figure 2B). Moreover, the CX and CMB cells maintained drug resistance *in vivo* when re-transplanted into mice and re-challenged with CX-5461 (Figure S2A) and CX-5461+EV (Figure S2B) respectively. EV cells showed little change in sensitivity to EV treatment, consistent with our previous finding that EV treatment did not provide a significant survival benefit in the Eμ-*Myc* B-lymphoma cells (clone #107) (*4*).

**Fig. 2:**
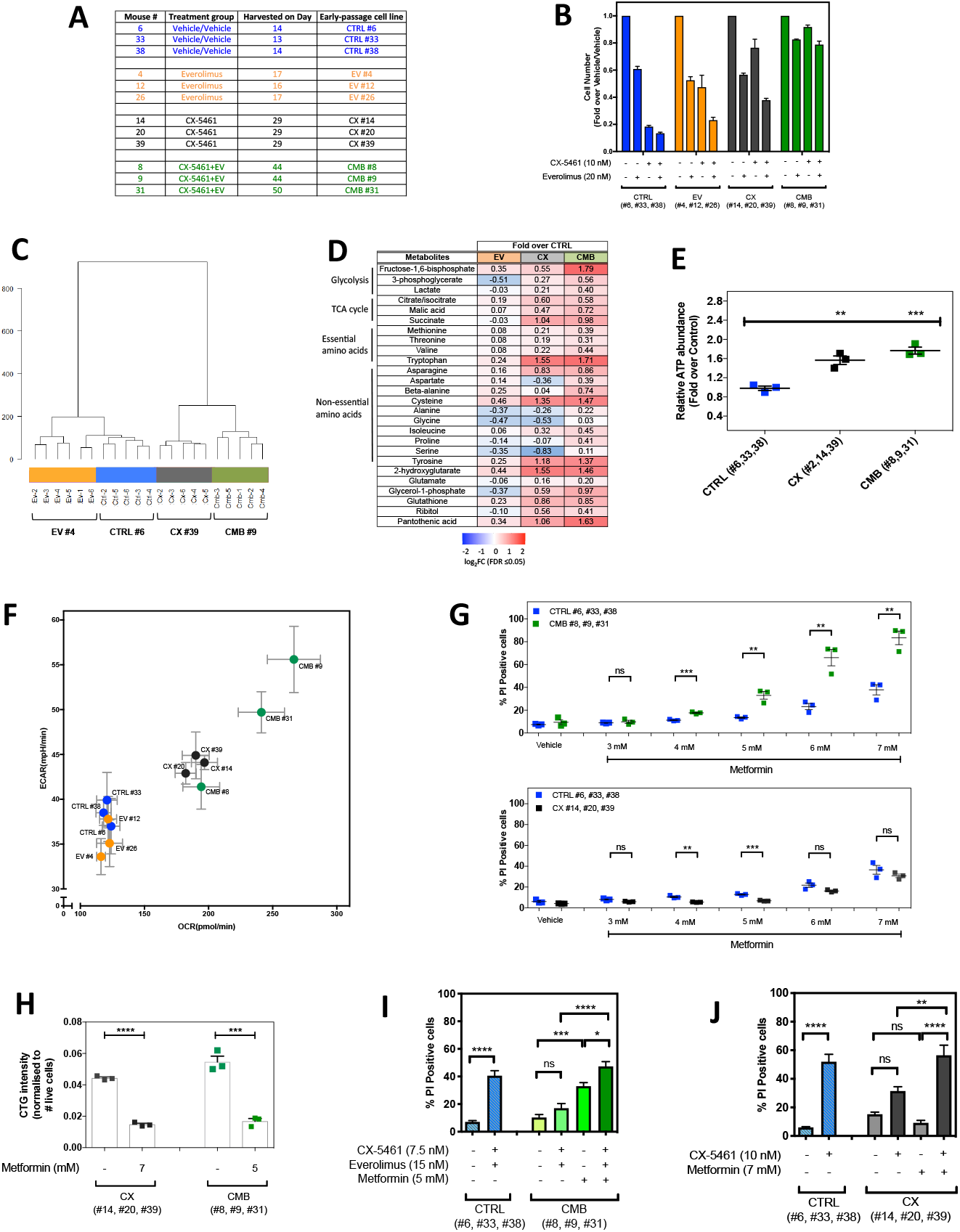
The early-passage Eμ-*Myc* B-lymphoma cells exhibit altered drug response and distinct metabolic profiles. **(A)** The early-passage Eμ-*Myc* B-lymphoma cells used in this study, which were established from mice that were drug-naïve (“CTRL” cell lines), previously treated with everolimus (“EV” cell lines) or CX-5461 (“CX” cell lines) alone, or combination of both (“CMB” cell lines). Harvest day indicates the number of days post-transplantation of lymphoma cells. **(B)** Cell viability analysis of the indicated early-passage Eμ-*Myc* B-lymphoma cells treated with indicated compound(s) for 24-hour as determined by Beckman® Coulter counter. Graphs represent mean ± SEM of n=3. **(C)** Hierarchical clustering analysis of metabolomics data (n=5-6) obtained from gas chromatography (GC)-mass spectrometry (MS) analysis of early-passage cell lines (n=5-6). **(D)** Steady-state abundance of indicated metabolites (fold-over CTRL cells; FDR ≤ 0.05) (n=5-6). **(E)** Intracellular ATP levels determined using the CellTiterGLO®-based assay in drug-naïve (CTRL) and CX-5461+EV-resistant (CMB) early-passage cell lines. Graphs represent mean ± SEM of n=3. **(F)** Basal extracellular acidification rate (ECAR) and oxygen consumption rate (OCR) in the early-passage cell lines determined using the Seahorse XF96 Extracellular Flux Analyzer; graphs represent mean ± SD of 6-8 technical replicates for each biological replicate (n=3). **(G)** Propidium iodide (PI) exclusion analysis of early-passage CX-5461-everolimus combination therapy-resistant (CMB), CX-5461-resistant (CX) and drug-naïve (CTRL) lymphoma cells treated with indicated concentrations of metformin for 48 hours. Graphs represent mean ± SEM of n=3. **(H)** CellTiterGLO®-based assay measuring cellular ATP levels of the CX-5461-resistant (CX) and CX-5461-everolimus combination therapy-resistant (CMB) cells treated with metformin as indicated for 48 hours. **(I)** PI analysis of CTRL and CMB cells treated with CX-5461and everolimus in the presence and absence of metformin for 48 hours as indicated. **(J)** PI analysis of CTRL and CX cells treated with CX-5461 in the presence and absence of metformin for 48 hours as indicated. Graphs represent mean ± SEM of n=3. (E), (G) & (H) Data were analyzed by Student’s t test. (I) & (J) Data were analyzed by one-way ANOVA. ns, not significant; *, P ≤ 0.05; **, P ≤ 0.01; ***, P ≤ 0.001; ****, P ≤ 0.001.

The early-passage cell lines maintained the appropriate on-target responses to these targeted therapies. Everolimus inhibited mTORC1 activity in all cell lines, as reflected by reduced phosphorylation of RPS6 (Ser240/244) (Figure S2C). As expected, CX-5461 treatment for 3 hours robustly decreased the rate of 47S pre-rRNA synthesis in the CTRL and EV cells (Figure S2D and S2E). Although the CX and CMB cells at 3 hours post-treatment were less sensitive to rDNA transcription inhibition (Figure S2F and S2G) compared to the CTRL and EV cells, longer term CX-5461 treatment resulted in robust on-target inhibition of Pol I transcription (Figure S2H-J). These findings were also consistent with the ability of CX-5461 to induce p53 accumulation in the cell lines (Figure S2K), confirming that loss of the IRBC is not associated with the mechanism of resistance to CX-5461 and CX-5461+EV.

Since acute targeting of cellular metabolism is associated with the cellular response to CX-5461+EV treatment (Figure 1E) and targeting metabolism improves the efficacy of ribosome-targeted therapy (Figure S1), we investigated whether metabolic re-wiring is associated with acquired resistance, by profiling metabolite levels in CTRL, EV, CX and CMB early-passage Eμ-Myc B-cell lymphoma cell lines using gas chromatography-mass spectrometry (GC-MS) (Figure 2C and 2D). Multivariate analysis clearly separated the metabolic phenotype of these cell lines (Figure 2C). Strikingly, the CX and CMB cell lines had elevated levels of glycolytic and tricarboxylic acid (TCA) cycle intermediates as well as a number of essential and non-essential amino acids compared to the EV and CTRL cell lines (Figure 2D). Consistent with our metabolomic analysis, the CX and CMB cells had elevated ATP content compared to the CTRL cells (Figure 2E). Seahorse XF96 analysis confirmed that CX and CMB cells were metabolically more active, as indicated by increased glycolysis and oxidative phosphorylation (Figure 2F). These results revealed energy and amino acid metabolism of CMB cells is indeed re-wired in response to the acute translation-dependent inhibition of metabolism when the drug-naïve cells were treated with CX-5461+EV (Figure 1E). Furthermore, while acute treatment with CX-5461 did not significantly affect metabolism (Figure 1E), consistent with the protective role of metformin-sensitive metabolism in limiting response of naïve cells to CX-5461 (Figure S1G), acquired resistance to CX-5461 was also associated with elevated cellular metabolism, particularly ATP production and oxidative phosphorylation (Figure 2E and 2F).

To test whether the re-wired metabolism in these CX and CMB cells contributes to drug resistance, we treated the CTRL, CX and CMB cells with metformin. CMB cells exhibited a greater sensitivity to metformin compared to the CTRL and CX cells (Figure 2G). Strikingly, metformin, at concentrations that resulted in similar levels of cell death and reduction in ATP abundance in both CX and CMB cells (Figure 2H), re-sensitized CMB cells to CX-5461+EV treatment, increasing the percentage of cell death to the levels achieved when CTRL cells were treated with CX-5461+EV (Figure 2I). Furthermore, metformin treatment increased CX-5461-induced cell death in CX cells (Figure 2J), consistent with the effects of metformin on both ATP levels (Fig 2H) and on mTORC1 signalling to P-RPS6 following activation of AMPK (Figure S2L). Thus, reprogramming of metformin-sensitive metabolism is integral for the acquired resistance to ribosome-targeting therapy.

### Increased translation of components of the cAMP pathway is associated with resistance to ribosome targeting therapy

To investigate whether there are translational alterations that underpin the energy metabolism-associated changes observed in cells resistant to ribosome-targeted therapy, we applied polysome profiling (Figure 1B) and performed pathway analysis using MetaCore® GeneGO on the mRNAs enriched on the polysomes of CMB cells to identify potential driver pathways of resistance. This analysis identified the cyclic-adenosine 3’,5’-monophosphate (cAMP) signaling pathway as the top hit when comparing CMB versus CTRL (Figure 3A), with increased translation efficiency of mRNAs encoding essential components of this pathway, including adenylate cyclase and cAMP-guanine nucleotide exchange factors (cAMP-GEFs) 1 and 2, which is also known as exchange protein directly activated by cAMP (EPAC) 1 and 2. Adenylate cyclase converts ATP into cAMP, which then activates protein kinase A (PKA) or EPAC1/2, the GEFs for activation of Ras-related protein 1 (RAP1), the two best characterised direct downstream targets of cAMP (*23, 24*). To confirm functional elevation of cAMP signaling, we performed liquid chromatography-MS (LC-MS), Western blot and pull-down analysis to demonstrate elevated cAMP abundance (Figure 3B), increased EPAC2 and GTP-bound RAP1 levels (Figure 3C), and polysome-association of *Rap1* mRNAs (Figure S3A) in the CMB cell lines compared to CTRL. In CX cells, cAMP signalling pathway was identified to be upregulated (Figure S3B), consistent with increased EPAC2 expression and GTP-bound RAP1 (Figure 3C) and polysome-association of *Rap1* mRNA (Figure S3A). Importantly, the expression levels of mRNAs from the genes encoding EPAC1 and 2 (*RAPGEF3* and *RAPGEF4)* are elevated in human hematological cancers including AML and diffuse large B-cell lymphoma (DLBCL) (Figures S3C and S3D). Indeed, EPAC1 and EPAC2 protein expression was elevated in the human AML cell lines MV4-11, SKM-1, SHI-1 and THP-1 compared to white blood cells from healthy patients (Figure S3E). Furthermore, using the EPAC1-specific inhibitor CE3F4 (*25*) and the EPAC2-specific inhibitor ESI-05 (*26, 27*), the viability of all four AML cell lines tested were reduced in response to EPAC1/2 inhibition (Figure S3G). Together, these data demonstrate the potential for a pro-survival role of the cAMP-EPAC1/2 pathway (*28*) in the etiology of hematological malignancies with deregulated MYC activity such as AML and B-cell lymphoma and in mediating cell survival in response of targeting the cell’s translational machinery.

**Fig. 3:**
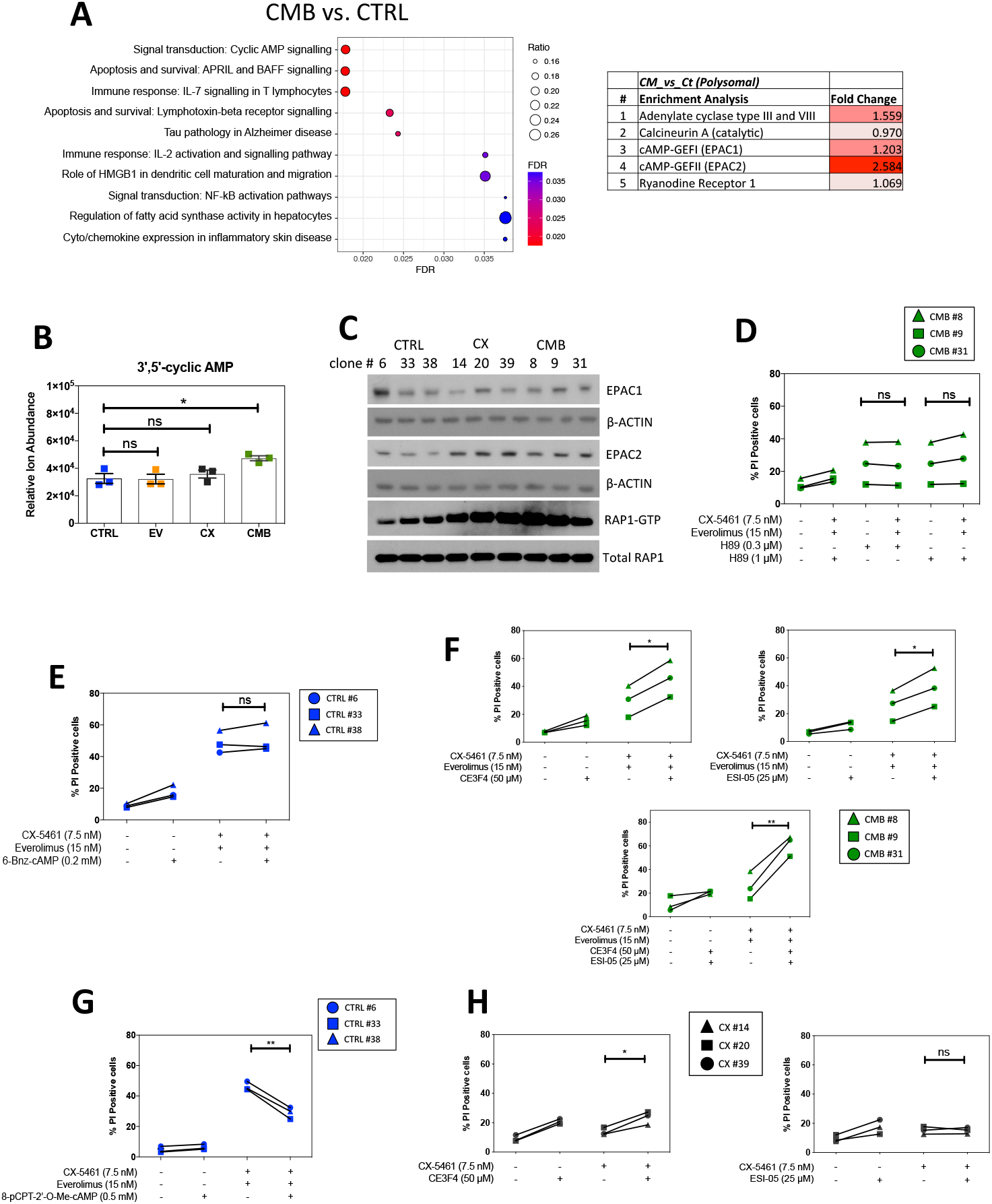
cAMP-dependent pathway mediates resistance to CX-5461-everolimus co-treatment. **(A)** Enrichment analysis by GeneGO MetaCore® of the polysomal RNAseq data comparing CX-5461+EV-resistant (CMB) and drug-naïve cells (CTRL) (n=3; false discovery rate (FDR) ≤ 0.05; fold change (FC) ≥ 1.5 or ≤-1.5). **(B)** Abundance of intracellular 3’5’-cyclic AMP as measured by a liquid chromatography (LC)-mass spectrometry (MS) analysis. Graphs represent mean ± SEM of n=3. Data were analyzed by one-way ANOVA. ns, not significant; *, P ≤ 0.05. **(C)** Western analysis of EPAC1 and EPAC2 abundance, as well as active GTP-bound RAP1 levels in the indicated early-passage cells during their log-phase growth period (n=3). Actin and total RAP1 were used as loading controls. **(D)** Propidium iodide (PI) exclusion assays of CMB cells treated with CX-5461 and everolimus as indicated in the presence or absence of a selective PKA inhibitor H89 for 24 hours. **(E)** PI exclusion analysis of early-passage drug-naïve (CTRL) lymphoma cells treated with CX-5461 and everolimus in the presence or absence of selective PKA activator 6-Bnz-cAMP. **(F)** PI exclusion analysis of the CMB cells treated with CX-5461 and everolimus as indicated in the presence or absence of EPAC1 inhibitor CE3F4 or EPAC2 inhibitor ESI-05 for 48 hours. **(G)** PI exclusion analysis of early-passage drug-naïve (CTRL) cells treated with CX-5461 and everolimus in the presence or absence of the selective EPAC activator 8-pCPT-2-O-Me-cAMP. **(H)** PI exclusion analysis of the CX cells treated with CX-5461 as indicated in the presence or absence of EPAC1 inhibitor CE3F4 or EPAC2 inhibitor ESI-05 for 48 hours. **(D, E, F, G)** Data were analyzed by paired one-way ANOVA. Graphs represent mean ± SEM of n=3. Black triangle: CX-5461-resistant (CX) cells clone #14, black square: CX cells clone #20, black circle: CX cells clone #39. Green triangle: CX-5461-everolimus combination therapy-resistant (CMB) cells clone #8; green square: CMB cells clone #9; green circle: CMB cells clone #31. Blue circle: early-passage drug-naive (CTRL) cells clone #6; blue square: CTRL cells clone #33; blue triangle: CTRL cells clone #38. ns, not significant; *, P ≤ 0.05; **, P ≤ 0.01.

To investigate whether altered cAMP signalling is fundamental for resistance to single-agent and/or combinatorial ribosome targeted therapy, we utilised specific inhibitors/activators of PKA and EPAC (*29, 30*). Inhibition of PKA by H89 (*29*) (Figure S3D) did not alter CMB cells’ response to CX-5461+EV treatment (Figure 3D). Similarly, 6-Bnz-cAMP, which robustly activates PKA signaling, as detected by a phospho-PKA substrate antibody (Figure S4B), was unable to protect CTRL cells from CX-5461+EV-induced cell death (Figure 3E) (*29, 31*). However, inhibition of EPAC1 and EPAC2 function, as measured by the reduced abundance of GTP-bound RAP1 (Figure S4C) by CE3F4 (*25*) and ESI-05 (*26, 27*) respectively, re-sensitized CMB cells to CX-5461+EV treatment (Figure 3F). Moreover, dual inhibition of EPAC1 and EPAC2 by combining CE3F4 and ESI-05 resulted in a more significant re-sensitization to CX-5461+EV treatment (Figure 3F), consistent with the proposed functional redundancy of the two isoforms (*32, 33*). These data indicate that EPAC1/2 activity, but not PKA, is required for the development of resistance to CX-5461+EV. Consistent with these findings, activation of EPAC1/2 in the CTRL cells using 8-pCPT-2’-O-Me-cAMP (*34, 35*), which increases RAP1-GTP abundance (Figure S4D), was able to reduce CX-5461+EV-mediated cell death (Figure 3G). This confirms that elevated EPAC1/2 activity contributes to protection from CX-5461+EV-mediated cell death and therefore the development of an energy- (Figure 2I) and cAMP-dependent resistant mechanism in the CX-5461+EV-resistant CMB cells. EPAC1/2 inhibitors did not re-sensitize CX cells to CX-5461 treatment (Figure S4E and Figure 3H) indicating that while the resistant mechanisms induced following single agent and combinatorial ribosome-directed treatments entail re-wired metabolism, there are subtle differences in the resulting requirements for cAMP signalling. This may be reflected in the lack of increased cAMP observed in CX vs CTRL cells (Figure 3B). In addition, comparison of the RNAseq data from polysomal mRNAs from CTRL, CX and CMB cell lines revealed that CMB cells upregulated the translation of mRNAs encoding multiple components of the mitochondrial electron transport chain (Figure S3F). These differences may explain CMB cells’ higher metformin sensitivity compared to CX cells (Figure 2G).

### CX-5461+EV-resistant cells are sensitized to metformin-mediated targeting of the cAMP-EPAC1/2-RAP1 pathway *in vivo*

The data presented above demonstrated that targeting oxidative phosphorylation or the cAMP-EPAC1/2-RAP1 signaling pathway re-sensitized the CMB cells to CX-5461+EV treatment. While there is currently no inhibitor of the cAMP-EPAC pathway in the clinic or in clinical trials, metformin is already being used in the clinic and can indeed reduce the intracellular abundance of cAMP (*36*). We hypothesized that metformin’s ability to sensitize CMB cells to CX-5461+EV treatment is associated with inhibition of cAMP-EPAC1/2-RAP1 signaling. The CMB and CX cells were treated with metformin as for Figure 2G and 2H. This treatment robustly lowered RAP1-GTP levels, but only in the CMB cells (Figure 4A), supporting a model (Figure 4B) where resistance to CX-5461+EV is, at least in part, mediated through cAMP-EPAC1/2-RAP1 pathway and can be suppressed using metformin (Figure 4A and Figures 2H-I).

**Fig. 4:**
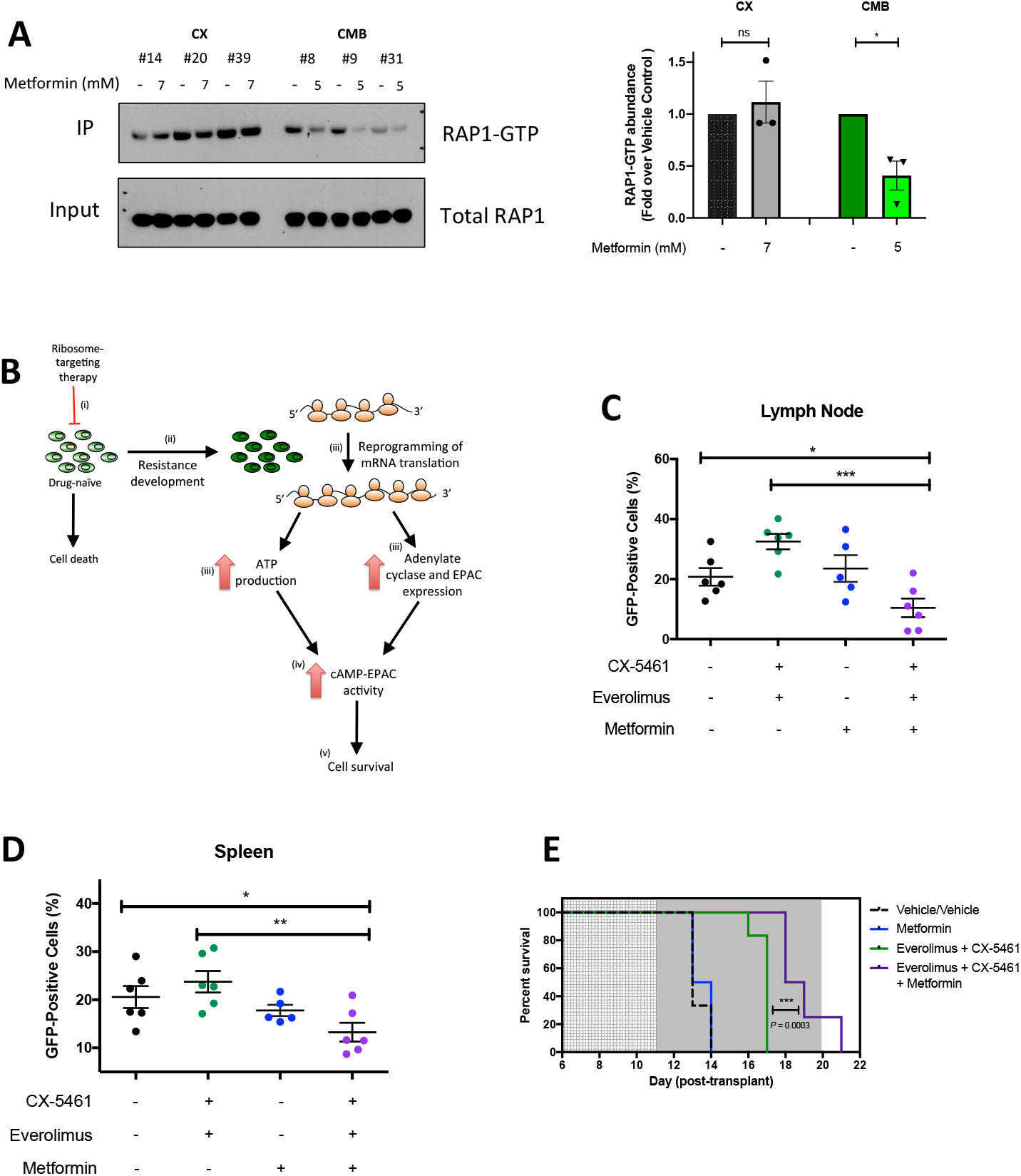
Metformin inhibits cAMP-EPAC1/2 signaling and re-sensitizes combination therapy-resistant early passage Eμ-*Myc* B-lymphoma cells to CX-5461-everolimus co-treatment *in vivo*. **(A)** Western analysis demonstrating the effects of metformin treatment for 48 hours on the levels of active GTP-bound RAP1 in CX and CMB cells (n=3) and its quantitation. **(B)** We proposed a model of translation-driven response to ribosome-targeting therapies where (i) drug therapy acutely inhibits the translation of key metabolic mRNAs (ii) continued compromised ribosome synthesis and translation allows selection of clones with elevated translation-dependent ribosome biogenesis; (iii) these clones exhibit specific upregulation of translation and synthesis of oxidative phosphorylation components (Figure S4F), EPAC and RAP1 proteins (Figures 3A, 3C and S3B); (iv) these in turn elevate ATP production (Figure 2E) and enhances its conversion into cAMP (Figure 3B); (v) cAMP is utilised by the EPAC proteins, thus protecting the resistant cells from activation of death mechanisms (Figures 3F-G) induced by CX-5461+EV co-treatment. Proportion of green fluorescent protein (GFP)-positive CMB (clone #8) cells in **(C)** lymph node and **(D)** spleen of transplanted C57BL/6 mice treated as indicated for 6 hours on day 12 post-transplant. Graphs represent mean ± SEM of 6 mice per group. **(E)** Kaplan-Meier curve of C57BL/6 mice transplanted with CX-5461-everolimus-resistant (CMB #8) early-passage Eμ-*Myc* B-lymphoma cells treated with vehicles (everolimus vehicle: 1% methylcellulose; CX-5461/metformin vehicle: 25 mM NaH_2_PO_4_); CX-5461 (35 mg/kg every twice weekly) and everolimus (5 mg/kg daily), metformin (400 mg/kg twice daily), or CX-5461, everolimus and metformin (35 mg/kg twice weekly, 5 mg/kg daily and 400 mg/kg twice daily, respectively). Data were analyzed by a log-rank (Mantel–Cox) test. Vehicle vs. CX-5461-everolimus: *P* = 0.0006, Vehicle vs. CX-5461-everolimus-metformin: *P* = 0.0001. CX-5461-everolimus vs. CX-5461-everolimus-metformin: *P* = 0.0003. (A, C, D) Graphs represent mean ± SEM of n=3. Data were analyzed by Student’s *t* test (A) or one-way ANOVA (C, D). ns, not significant; *, P ≤ 0.05; **, P ≤ 0.01; ***, P ≤ 0.001; ****, P ≤ 0.0001.

To investigate whether metformin could also re-sensitize the CMB cells to CX-5461+EV treatment *in vivo*. CMB lymphoma-bearing C57BL/6 mice were pre-treated with metformin (*37*) or vehicle for 3 days (600 mg/kg twice daily on the first and second day; 500 mg/kg twice daily on the third day) and then treated for 6 hours with CX-5461 and EV as single agents or in combination. As expected, CX-5461+EV did not alter the abundance of GFP-positive drug-resistant CMB cells in the lymph node (Figure 4C) or spleen (Figure 4D) compared to vehicle pre-treated mice. Importantly, the single-agent metformin had no effect on the number of GFP-positive cells, whereas they were significantly reduced in the lymph nodes (Figure 4C) and spleens (Figure 4D) of mice receiving the CX-5461+EV plus metformin treatment. Thus, metformin was able to re-sensitize the CMB cells to CX-5461+EV treatment *in vivo*.

To determine whether the triple combination therapy could enhance survival of mice bearing this highly aggressive disease, C57BL/6 mice transplanted with the CMB cells were pre-treated for 5 days with metformin (500 mg/kg twice daily) then at Day 11 post-transplant, treated with either: (i) vehicle; (ii) CX-5461+EV (35 mg/kg every Mon-Wed-Fri and 5 mg/kg daily, respectively); iii) metformin (400 mg/kg twice daily); or (iv) CX-5461+EV+metformin triple combination therapy. The triple combination significantly increased the survival window compared to all the groups including the CX-5461+EV treatment (Figure 4E). Together these data show that the re-sensitization of the highly aggressive, drug-resistant CMB cells to CX-5461+EV treatment in the presence of metformin *in vitro* was reproducible *in vivo,* thus providing proof-of-principle that combined inhibition of translational machinery and energy metabolism for the treatment of highly aggressive MYC-driven lymphoma can provide a therapeutic window.

## Discussion

Oncogene-driven cancers are characterized by elevated ribosome biogenesis that can be targeted with specific inhibitors of ribosomal RNA synthesis to treat refractory blood cancer. We reported that CX-5461 potently enhances the efficacy of PI3K/AKT/mTORC1 inhibitors in treating MYC-driven lymphoma (*4*). Here we use a combination of polysome profiling and metabolomics analysis of tumors from acutely treated mice and early-passage cells from acquired-resistant tumors to interrogate the mechanisms underpinning this ribosome-directed therapeutic approach.

We demonstrate that the synergistic efficacy of combining CX-5461 with the mTORC1 inhibitor everolimus is associated with dramatic suppression of the translation of mRNAs encoding key proteins controlling mRNA translation itself. This specific and acute targeting of the ribosome *in vivo* leads to selective inhibition of the translation of mRNAs containing longer 5’UTRs (Figure S1D), including those encoding proteins critical for the regulation of energy metabolism, with the subsequent result being increased lymphoma cell death. This is distinct from a report that mTOR regulated the translation of mRNAs containing shorter, less complex 5’UTRs (*14*), suggesting that the translational changes we observed in response to CX-5461+EV treatment are not solely driven by EV-dependent inhibition of mTORC1 signaling, but a specific result of the combination therapy.

Since p53-null Eμ-*Myc* B-cell lymphomas are resistant to CX-5461+EV treatment (*4*) we initially postulated that mutation or loss of p53 might underlie the resistance mechanism. Indeed, loss of *TP53* is observed in ~40% of human Burkitt lymphoma cases (*38, 39*). However, we observed that the p53 responses were maintained in acquired resistant cells. Instead, our studies define a new model of molecular response of oncogene-driven cancer to ribosome-targeted therapy (Figure 4B).

Under the pressure of the combination treatment, tumor cells undergo translational re-programing, which resulted in increased translation of polysome-associated mRNAs that encode components of the mitochondrial respiratory electron transport chain and the cAMP-EPAC1/2-RAP1 pathway. The resulting increase in ATP and cAMP production (Figures 2E and 3B) further activates EPAC signaling to RAP1, an energy-dependent pro-survival pathway that provides protection from drug induced cell death (*28*). Functionally, specific inhibition of cAMP-EPAC1/2-RAP1 signaling re-sensitizes resistant cells to combination therapy demonstrating that this pro-survival pathway is a critical mediator of resistance (Figures 3D-G). Importantly, targeting elevated energy metabolism with the anti-diabetic drug metformin sensitized both CX cells to CX-5461 and CMB cells to CX-5461+EV. In CX cells, drug-resensitization by metformin was associated with AMPK-mediated inhibition of mTORC1 (Fig 2H and S2L). In CMB cells, metformin was able to inhibit EPAC1/2-RAP1 signaling and markedly improved the efficacy of combination therapy in resistant disease *in vitro* and *in vivo* (Figures 4A, 4C, 4D and 4E). Moreover, we highlighted the association of elevated EPAC1/2-RAP1 signaling with poor prognosis in AML and DLBCL, as well as the potential of improving CX-5461 treatment in AML. Our findings demonstrate that elevated metabolic activity and energy production is not merely a hallmark of cancer associated with uncontrolled growth, but can be harnessed to activate the pro-survival signaling through cAMP-EPAC1/2-RAP1, a new metabolic vulnerability that can be exploited to further improve the efficacy of ribosome-targeting therapy. This makes a convincing case for the importance of developing inhibitors to EPAC function with improved pharmacological properties as an additional tool for targeting oncogene-driven cancer. Thus, we believe this approach of co-targeting metabolism and Pol I-directed ribosome targeting therapy provides a new paradigm for improving the efficacy of metabolic cancer therapies, as well as both traditional chemotherapeutics that target ribosome biogenesis and the newly developed low genotoxic approaches including CX-5461 and BMH-21 (*1*).

Finally, this study reinforces the recent finding that genetically compromised ribosome biogenesis results in specific re-wiring of translation that underlies impaired erythroid differentiation (*15*). More broadly, diseases of genetically compromised ribosome biogenesis such as Diamond Blackfan anaemia, are similarly characterized by specific alterations in mRNA translation (*15*) and by significantly increased incidence of cancer, a paradox termed “Dameshek’s riddle” (*40*). This riddle is reinforced by observations of somatically acquired mutations and deletions in ribosomal proteins in T-cell acute lymphoblastic leukemia as well as solid tumors, such as gastric and ovarian cancer (*40–43*). Despite intense investigation, the mechanisms by which genetically compromised ribosome biogenesis leads to increased cancer susceptibility in patients with ribosomopathies remain a mystery. Our findings raise the possibility that at least part of the answer to Dameshek’s riddle lies in specific rewiring of translation in response to chronically compromised ribosome biogenesis, whereby the subsequent translationally-driven elevated metabolism and pro-survival mechanisms promote malignant transformation later in life.

## Methods

### Cell culture and reagents

Eμ-*Myc* B-lymphoma cells (MSCV *Gfp)* were cultured in Anne Kelso DMEM, supplemented with 10% heat-inactivated (h.i.) fetal bovine serum (FBS), 100 μM L-asparagine (Merck), penicillin/streptomycin/glutamine (Thermo Fisher Scientific) and 0.5% beta-mercaptoethanol. Early-passage Eμ-*Myc* B-lymphoma cells were established by processing lymph nodes isolated at the end of a survival experiment in (*4*). These early-passage Eμ-*Myc* B-lymphoma cells were cultured in Anne Kelso DMEM supplemented with 20% h.i. FBS, 100 μM L-asparagine (Merck), penicillin/streptomycin/glutamine (Thermo Fisher Scientific) and 0.5% beta-mercaptoethanol. Human acute myeloid lymphoma cell lines (MV4-11 (MLL-AF4; p53 wild-type (WT), SKM-1 (EZH2; p53-mutant (R248Q)), SHI-1 (MLL-AF6; p53WT), THP-1 (t(9;11); p53-null R174fs)) were cultured in RPMI 1640 medium plus HEPES supplemented with 20% h.i. FBS, 4 mM Glutamax (Thermo Fisher Scientific) and 1% v/v antibiotics/antimycotics (Thermo Fisher Scientific). Everolimus (S1120) was purchased from Selleckchem. CX-5461 was purchased from SYNkinase (SYN-3031). Metformin was purchased from Merck (D150959). CEF34 was purchased from Cayman Chemicals (17767). ESI-05 and 8-pCPT-2-O-Me-cAMP were purchased from Biolog (catalogue number M092 and C041, respectively). 6-Bnz-cAMP was purchased from Tocris Bioscience (5255). H89 was purchased from Merck (B1427).

### Animal experiments

All animal experiments were performed with approval from the Animal Experimentation Ethics Committee at the Peter MacCallum Cancer Centre (Ethics number E462 and E557). For *in vivo* drug studies, 2 × 10^5^ Eμ-*Myc* lymphoma cells were injected into the tail vein of 6-8 week old male C57BL/6 mice (Walter and Eliza Hall Institute, Australia). Disease onset/progression was monitored by peripheral blood (tail bleed) analysis for green fluorescent protein (GFP). Mice were treated with pharmacologic inhibitors from 10 days post-transplant (survival studies) or on day 12 post-transplants (acute studies). Everolimus was administered daily at 5 mg/kg via oral gavage in 5% dimethyl sulfoxide (DMSO) in 1% methylcellulose. CX-5461 was administered twice weekly (unless otherwise indicated in figure legends) at 35 mg/kg via oral gavage in 25 mM NaH_2_PO_4_ (pH 4.5). Metformin was administrated twice daily (2 × 500 mg/kg unless otherwise indicated in figure legends) via oral gavage in 25 mM NaH2PO4 (pH 4.5). For survival studies, mice were dosed until an ethical end-point was reached (enlarged lymph nodes, inactivity, hunched posture, laboured breathing, weight loss (equal to 20% of initial body weight) and ruffled fur). White blood cells, lymph node cells, and spleen cells were analyzed for GFP and B220 (CD45R) (Thermo Fisher Scientific 17-0452-82) expression using BD CantoII or BD Fortessa. Flow cytometry data was analyzed with Flowlogic software (Inivai Technologies). Snap-frozen lymph nodes or spleens were homogenized using a Precellys 24/Cryolys cryomill (Bertin Technologies) (6,800 rpm; 2 × 30-second pulse, 45 seconds interval between pulses; 0 **°**C).

### Protein analysis and Western blotting

Protein was extracted with SDS-lysis buffer (0.5 mM EDTA, 20 mM HEPES, 2% (w/v) SDS pH 7.9) and protein concentrations were determined with the Bio-Rad DC protein assay. Proteins were resolved by SDS-PAGE, transferred to PVDF membranes and immunoblotted with primary and horseradish peroxidase–conjugated secondary antibodies (Supplementary Table). Protein was visualized by Amersham enhanced chemiluminescence (ECL) Western Blotting Detection Reagent (GE Healthcare Life Sciences) and X-ray film (Fujifilm SuperRX). Active RAP1 pull-down experiments were performed using the RAP1 Activation Assay (Abcam ab212011) kit according to manufacturer’s instructions.

### Polysome profiling, RNA isolation and RNAseq analysis

Suspension cells were harvested by removing the culture media following centrifugation (400 g, 4 minutes, 4 **°**C). The cells were washed with ice-cold hypotonic wash buffer (5 mM Tris pH 7.5; 1.5 mM KCl; 2.5 mM MgCl2 and 100 μg/mL cycloheximide (Sigma)) and the supernatant was removed by centrifugation (400 g, 4 minutes, 4 **°**C). A hypotonic lysis buffer (5 mM Tris pH 7.5; 1.5 mM KCl; 2.5 mM MgCl_2_; 0.5% Triton-X; 0.5% sodium deoxycholate; 1X EDTA-free protease inhibitor; 2 mM DTT; 10 μL RNAsin (Promega); 100 μg/mL cycloheximide) was added to the cell pellet. The samples were mixed by pipetting and centrifuged (16,000 g, 7 minutes, 4 **°**C). The supernatant (cytoplasmic lysate) was transferred to a fresh microfuge tube. 10% of the cytoplasmic lysate was transferred into a microfuge tube containing 500 μL TRIzol® reagent. The remaining lysate was layered on top of a linear 10-40% sucrose gradient, ultracentrifuged (36,000 rpm, 2¼ hours, 4 **°**C) and fractionated using the Foxy Jr Fraction Collector with constant monitoring of absorbance at 260 nm by an ISCO UA-6 Absorbance Detector (Teledyne). Cytoplasmic (input) RNAs and polysomal RNAs from pooled fractions corresponding to four or more ribosomes were isolated using a TRIzol®-based method and purified using the Qiagen RNeasy® Minikit according to manufacturer’s instructions, and analysed by RNAseq. RNA concentration was determined using a NanoDrop Spectrophotometer (Thermo Scientific) and integrity was evaluated using the RNA Nano Kit and 2100 bioanalyzer (Agilent Technologies).

cDNA libraries were generated using a ribodepletion method (TruSeq Ribo Profile Mammalian Kit, Illumina). Fragment sizes were evaluated using the DNA 1000 kit and 2100 bioanalyzer (Agilent Technologies). Libraries were subjected to single-end sequencing (SE50bp) (Hiseq2500, Illumina) to generate ~30 million reads per sample using standard protocols. Reads were aligned with Tophat2 and reads counts were obtained with htseq with the mouse genome (Ensembl release 78) as a reference, using an in-house bioinformatics pipeline, seqliner (http://bioinformatics.petermac.org/seqliner/). Differential translation was detected using anota2seq analysis (*16*) of polysome-associate mRNAs, using the total amount of cytosolic mRNA as a reference. Only coding genes with a count of at least 1 in any of the samples were considered. Samples were normalized using the “voom” option in anota2seq. An omnibus analysis was first performed in anota2seq to enrich for genes affected by any of the treatments, using a groupRvmPAdj cutoff of 0.15. Differential translation analysis was subsequently performed using anota2seq on this reduced set. Only genes with an apvRvm adjusted p-value < 0.05 and an apvEff (fold change) greater than 1.25 were considered to be significant. To perform differential analysis of polysome data in resistant studies (i.e. to compare mRNAs that were differentially bound to polysomes of treated vs. control cells), we used LIMMA version 3.28.21 (Ritchie et al., 2015) from Bioconductor. Only coding genes with an average logCPM > 1 were considered. Genes were considered to be significant if their adjusted p-values after Benjamini & Hochberg correction were less than 0.05, and their fold change was greater than 1.5.

For *in vivo* translational profiling experiments, cytoplasmic lysate was obtained by grinding snap-frozen tissue samples in liquid nitrogen proof containers submerged in dry ice-100% ethanol slurry to a fine powder. A modified lysis buffer (10-fold higher concentration of cycloheximide compared to the one used for cultured cells; RNAsin was replaced with 10 mM ribonucleoside vanadyl complex (New England Biolabs S1402S) was added to the ground tissue. The sample was transferred to an ice-cold dounce homogeniser and homogenised (30-60 strokes). The homogenate was transferred to a chilled microfuge tube and centrifuged (16,000 g, 7 minutes, 4 °C). The supernatant (cytoplasmic lysate) was transferred to a fresh microfuge tube. 10% of the cytoplasmic lysate was transferred into a microfuge containing 500 μL TRIzol® reagent. The remaining lysate was loaded onto linear (10-40%) sucrose gradients and processed as described above.

For the 5’UTR analysis, 5’UTR sequences were obtained using the biomaRt and org.Mm.eg.db R packages from the Mouse Genome GRCm38.p6. Only 5’UTRs shorter than 500 nucleotides were considered, since others have found that longer 5’UTRs were likely to be false positives (*15*). In cases where there were multiple 5’UTR for a single gene, we only considered the shortest 5’UTR. For the resistant study, only genes with FC > 1.5 or < −1.5 were considered, with adjusted *P* < 0.05. Only expressed genes were used as background. For the acute study, metabolic downregulated genes were those with an appEff (fold change) < 0, and an apvRvmPAdj (adjusted *P*) < 0.1 (from the omnibus-sorted genes). The minimum free energy of 5’UTR sequences was calculated using the ViennaRNA package version 2.4.7. In cases where there were multiple 5’UTR for a single gene, we only considered the 5’UTR with the lowest minimum free energy. One-sided Wilcoxon tests were used to determine significance.

### Cell viability assay

Cells were seeded in 96-well plates. 24 hours later, cells were treated with pharmacological compounds as indicated in the figure legends or text. Cells were stained with 1 μg/mL PI (Merck P4170) and analyzed using the FACSVerse (BD Biosciences). Flow cytometry data was analyzed with Flowlogic software (Inivai Technologies). Cell number was determined using the Z2 Coulter Counter (Beckman Coulter 383550) or CellTiterGLO®-based assay followed by luminescence reading by Cytation 3 Cell Imaging Multimode Reader (BioTek).

### ^32^P-orthophospate labeling

Eμ-*Myc* B-lymphoma cells (3 × 10^6^) were cultured in 3 ml media in the presence of 0.5 mCi of 32P-orthophosphate for 15 min. Cells were harvested on ice and RNA extracted using the QIAGEN RNeasy Minikit according to the manufacturer’s instructions. RNA (5 μg) was run overnight on a 1.2% MOPS/formaldehyde agarose gel. The gel was dried using a Model 583 Gel Drier (Bio-Rad), exposed overnight to a phospho-imager screen (Molecular Dynamics) and scanned using a Typhoon Trio Variable Mode Imager (GE Healthcare). Band intensities were quantitated using Image Quant TL software (GE Healthcare).

### Bioenergetics analysis using the Seahorse XF96 Extracellular Flux Analyzer

All bioenergetics analyses were performed using the Seahorse Bioscience XF96 extracellular flux analyser (Seahorse Bioscience, Billerica, USA). Cells were washed with the assay media (unbuffered DMEM, 5 g/L glucose, 5 mM glutamine, 1 mM sodium pyruvate), before seeding in Seahorse XF96 96-well plates coated with Cell-Tek (3.5 ug/cm2, Corning) at 2 × 10^5^ cells/well in 180 μl of the assay media. The plate was centrifuged at 400g for 5 minutes at room temperature to allow the cells to form a monolayer. The plate was equilibrated in a non-CO2 incubator for 30 minutes prior to assay. The assay protocol consisted of 3 repeated cycles of 3 minutes mixing and 3 minutes of measurement periods, with oxygen consumption rate (OCR) and extracellular acidification rate (ECAR) determined simultaneously. Basal energetics were established after three of these initial cycles, followed by exposure to the ATP synthase inhibitor, oligomycin (1 μM) for three cycles, then p-trifluoromethoxy-phenylhydrazone (FCCP, 1.5 μM), which uncouples oxygen consumption from ATP production, was added for a further three cycles. Finally, the mitochondrial complex III inhibitor antimycin A (0.5 μM) and the complex I inhibitor rotenone (0.5 μM) was added for three cycles. At the completion of each assay, the cells were stained with 10 μM Hoechst. Images were analysed using a Cellomics Cellinsight 1 to determine the cell number per well.

### Gas chromatography – mass spectrometry (GC-MS)

Cells were quenched by transferring cell suspensions (5 mL) into 50 mL falcon tubes containing 35 mL ice-cold saline, followed by centrifugation (500 × g, 3 minutes, 0 **°**C). Once the supernatant was removed, the cells were re-suspended in 1 mL ice-cold saline, transferred to 1.5 mL microfuge tubes, and centrifuged (10,000 × g, 30 seconds, 0 **°**C). Supernatant was removed completely and metabolites were immediately extracted in 250 μL chloroform:methanol:water (CHCl_3_:CH_3_OH:H_2_O, 1:3:1 v/v; monophasic mixture) containing 0.5 nmoles scyllo-inositol as internal standard. The extracts were vortexed (~10 seconds) and incubated on ice for 15 minutes with additional vortexing every 5 minutes. Insoluble material was removed by centrifugation (10,000 × g, 5 minutes, 0 **°**C). H_2_O was then added to the supernatant to adjust the ratio of CHCl_3_:CH_3_OH:H_2_O to 1:3:3 (v/v; biphasic partition). The supernatant was vortexed and centrifuged (10,000 × g, 5 min, 0 **°**C) to induce phase separation. The upper (polar) phase was transferred to a fresh microfuge and dried *in vacuo* (30 **°**C) in 250 μL glass vial inserts. Samples were analyzed by GC-MS as previously described (*44*). Briefly, samples were prepared for GC-MS analysis using a Gerstel MPS2 autosampler robot. Polar metabolites were first methoximated with methoxyamine hydrochloride in pyridine (Sigma, 20 μl, 30 mg/ml, 2 hours, 37 **°**C) and then derivatized with TMS reagent (N,O-bis(trimethylsilyl) trifluoroacetamide containing 1% trimethylchlorosilane (Pierce, 20 μl, 1 hour, 37 **°**C) and analysed by GC-MS using a VF5 capillary column (Agilent, 30 m, 250 μm inner diameter, 0.25 μm film thickness), with a 10 m inert eziguard. The injector insert and GC-MS transfer line temperatures were 250 and 280 **°**C, respectively. The oven temperature gradient was programmed as follows: 35 **°**C (1 minute); then 25 **°**C /minute to 320 **°**C; and held at 320 **°**C for 5 minutes. Metabolites were detected by mass selective detector following electron ionization (−70 eV), where the scan range was 50-600 amu at 9.2 scans/sec. Metabolites were identified and areas integrated based on GC retention time and mass spectra as compared with authentic standards and in conjunction with MSD ChemStation Data Analysis Application (Agilent) using in-house and Wiley metabolite libraries. Statistical analyses were performed using Student *t* test following log transformation and median normalization. Metabolites were considered to be significant if their adjusted p-values after Benjamini-Hochberg correction were less than 0.05.

### Liquid chromatography – mass spectrometry (LC-MS)

Cells were quenched by transferring cell suspensions (5 mL) into 50 mL falcon tubes containing 35 mL ice-cold saline, followed by centrifugation (500 × g, 3 minutes, 0 **°**C). Once the supernatant was removed, the cells were re-suspended in 1 mL ice-cold saline, transferred to 1.5 mL microfuge tubes, and centrifuged (10,000 × g, 30 seconds, 0 **°**C). Supernatant was removed completely and metabolites were immediately extracted using ice-cold 80% acetonitrile:water (v/v) containing internal standards (^13^C-Sorbitol and ^13^C-Valine; final concentration 4 μM both). Metabolites in the cell extracts were analyzed by high-performance liquid chromatography mass spectrometer (LC-MS) as previously described (*44*). Briefly, metabolites were separated on an Agilent Technologies 1200 series HPLC system (Agilent Technologies, Santa Clara, US, USA) using a SeQuant ZIC–pHILIC (5 μm polymer) PEEK 150 × 4.6 mm metal-free HPLC column maintained at 25°C with solvent A (20 mM (NH_4_)_2_CO_3_, pH 9.0; Sigma-Aldrich) and solvent B (100% acetonitrile) at a flow rate of 300 μl/minute. Metabolites were detected by mass spectrometry on an Agilent Technologies 6545A series quadrupole time of flight mass spectrometer (QTOF MS) using an electrospray ionization source (ESI). LC-MS data was collected in negative MS mode. Data processing was performed using Agilent MassHunter workstation quantitative analysis for TOF software (Version B.07.00/Build 7.0.457.0). Metabolite identification was based on accurate mass, retention time and authentic chemical standards at level 1 confidence according to the Fiehn’s Metabolomics Standard Initiative.

## Data Analysis

Statistical tests were performed as described in Figure legends with GraphPad Prism Software (Version 7), or as outlined in relevant polysome profiling or metabolomics methods sections detailing their respective bioinformatics analysis. In AML studies, visualization of expression analysis was generated using the gene expression profiling interactive analysis (GEPIA) bioinformatics tool (*45*).

## General

The authors thank Kerry Ardley, Susan Jackson and Rachael Walker for technical assistance with animal experiments. They also thank the Peter MacCallum Cancer Centre Animal Facility, Molecular Genomics Core, FACS Facility, Laboratory Services and Media Kitchen. The authors would like to acknowledge Dr Maurits Evers (ANU) for his input into the visualization of the next-generation sequencing data.

## Funding

This work was supported by project grants and fellowships (M.J.M., R.D.H., G.A.M and R.B.P.) and a program grant (R.D.H., G.A.M and R.B.P.) from the National Health and Medical Research Council (NHMRC) of Australia. E.K. was supported by a University of Melbourne International Fee Remission Research Scholarship (MIFRS), a Melbourne International Research Scholarship (MIRS) and a Cancer Therapeutics (CTx) PhD Top Up Scholarship. A.S.T was supported by a University of Melbourne MIFRS and a Melbourne International Engagement Award (MIEA).

## Author contributions

Conception and design: E.P.K, R.D.H., K.M.H., J.K. and R.B.P.. Acquisition of data: E.P.K., J.R.D., D.P.D and J.K. Analysis and interpretation of data: E.P.K, A.S.T, D.L.G., O.L., M.J.M, K.M.H., J.K. and R.B.P. *In vivo* animal experiments were carried out with the assistance of C.C. and her team. Administrative, technical or material support: J.R.D, K.T.C, D.P.D, G.A.M., R.D.H. and E.S. Writing and revision of the manuscript: E.P.K., G.T., K.M.H., J.K. and R.B.P. Study supervision: K.M.H., J.K. and R.B.P.

## Competing interests

G.A. McArthur reports receiving commercial research grants from Celgene, Novartis, and Ventana and is a consultant/advisory board member for Provectus. R.D. Hannan is a consultant/advisory board member for Pimera, Inc. No potential conflicts of interest were disclosed by the other authors

## Notes

#### Summary of Updates

Supplemental Files updated.

