## Supplemental Figures S1 to S4 for "Reprogrammed mRNA translation drives resistance to therapeutic targeting of ribosome biogenesis"

**Affiliations**
[1] Peter MacCallum Cancer Centre, 305 Grattan St, Melbourne, Victoria, Australia, 3000.
[2] Sir Peter MacCallum Department of Oncology, The University of Melbourne, Parkville, Victoria, Australia, 3010.
[3] Department of Oncology-Pathology, Science for Life Laboratory, Karolinska Institutet, 171 65 Solna, Sweden.
[4] Metabolomics Australia, Bio21 Molecular Science and Biotechnology Institute, Parkville, Victoria, Australia, 3010.
[5] Department of Biochemistry and Molecular Biology, The University of Melbourne, Parkville, Victoria, Australia, 3010.
[6] Metabolism and Cancer Group, Molecular Mechanisms and Experimental Therapy In Oncology Program, Bellvitge Biomedical Research Institute, IDIBELL, 08908 Barcelona, Spain
[7] Department of Clinical Pathology, The University of Melbourne, Parkville, Victoria, Australia, 3010.
[8] ACRF Department of Cancer Biology and Therapeutics, The John Curtin School of Medical Research, ACT, Australia, 2601.
[9] Department of Biochemistry and Molecular Biology, Monash University, Clayton, Victoria, Australia, 3168.
[10] School of Biomedical Sciences, University of Queensland, Brisbane, Queensland, Australia, 4072.
† These authors contributed equally

### Supplementary Materials:

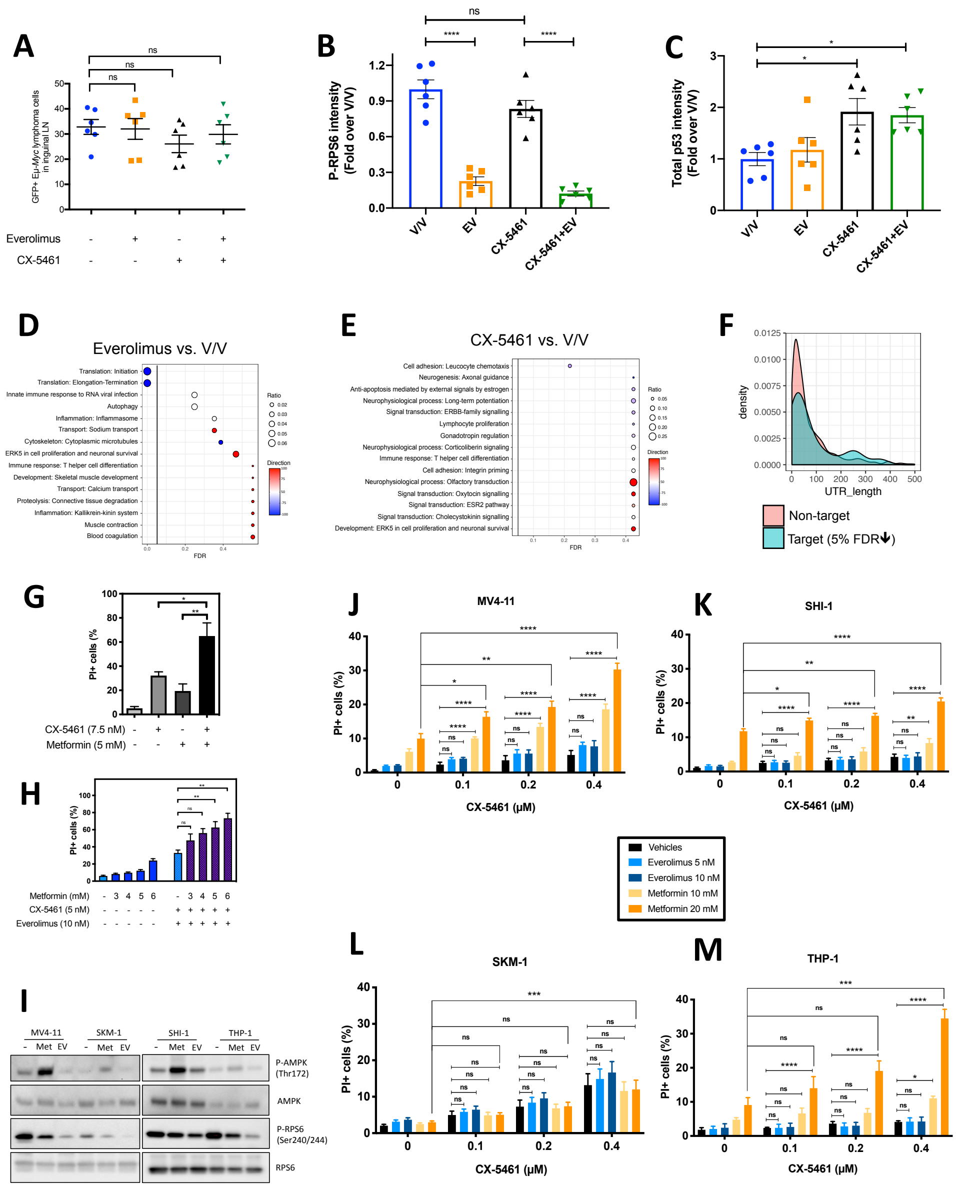


**Fig. S1. *In vivo* transcriptome-wide analysis of cellular response to ribosome targeting therapy**

**(A)** Flow cytometry analysis of inguinal lymph node cells isolated from C57BL/6 mice with transplanted Eμ-Myc B-cell lymphoma cells treated as indicated for 2 hours. 10,000 viable cells of the correct morphology were collected for each sample. Graphs represent mean ± SEM of n=6. Quantitation of **(B)** phosphorylated RPS6 levels normalised to total RPS6 and **(C)** p53 protein levels normalised to Actin in Figure 1A. Graphs represent mean ± SEM of n=6. **(D and E)** Enrichment analysis by MetaCore® GeneGO of genes in "translation up" and "translation down" categories identified by anota2seq analysis comparing lymph node cells isolated from mice in **(D)** CX-5461 single-agent treatment group or **(E)** everolimus single agent treatment group with those isolated from mice in vehicle group (n=6). "Ratio" is the value obtained by dividing the number of genes in an indicated molecular process that is found in our data with the number of genes curated in MetaCore®'s database. "Direction" visualises the percentage of genes associated with indicated pathways that are up (red) or down (blue). **(F)** Length of the 5'UTRs of metabolism-associated mRNAs that were downregulated in response to CX-5461+EV treatment *in vivo* (n=6). Propidium iodide exclusion analysis of Eμ-Myc B-lymphoma cells treated with **(G)** CX-5461 treatment in the presence and absence of metformin, or **(H)** CX-5461+EV treatment in the presence and absence of metformin as indicated for 48 hours. **(G and H)** Graphs represent mean ± SEM of n=3 independent experiments and data were analysed by one-way ANOVA. **(I)** Western blot analysis of phosphorylated AMPK and phosphorylated RPS6 for the on-target effect of 10 mM metformin treatment for 24 hours and 5 nM everolimus treatment for 3 hours respectively. Propidium iodide (PI) exclusion analysis of **(J)** MV4-11, **(K)** SHI-1, **(L)** SKM-1, **(M)** THP-1 human acute myeloid leukemia cell lines in response to treatments with CX-5461, everolimus (EV) and metformin as indicated for 72 hours. Data were analysed by two-way ANOVA. ns, not significant; P≥ 0.05; *, P ≤ 0.05; **, P ≤ 0.01; ***, P ≤ 0.001; ****, P ≤ 0.0001.

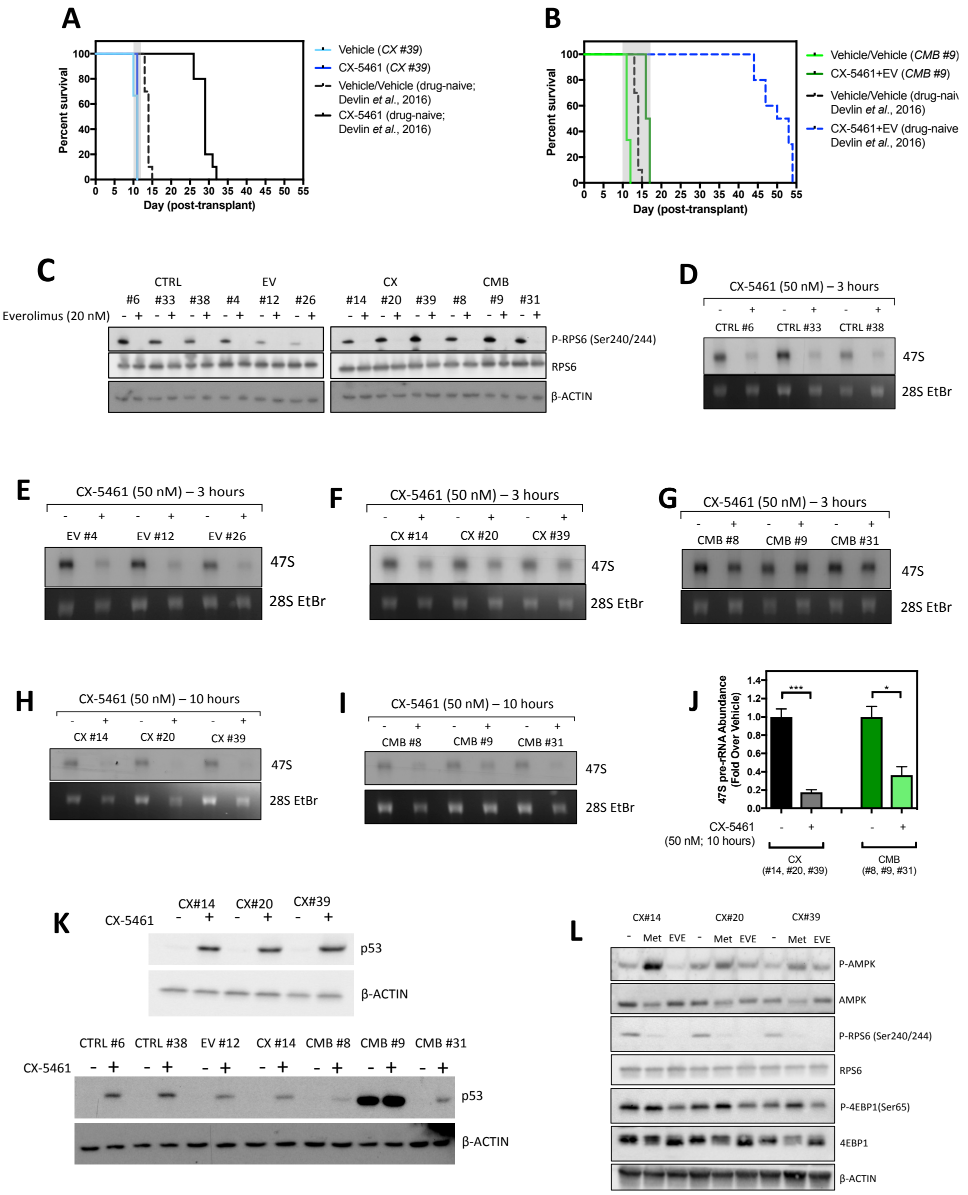


**Fig. S2. Everolimus and CX-5461 maintained their appropriate on-target effects in the early-passage Eμ-Myc lymphoma cell lines**

**(A)** Kaplan-Meier curve of C57BL/6 mice transplanted with the CX-5461-resistant CX #39 early-passage Eμ-*Myc* B-lymphoma cells treated with CX-5461 (35 mg/kg thrice weekly) (n=5-6). Survival curves of drug-naïve cells were actual data reproduced from Devlin et al., 2016. **(B)** Kaplan-Meier curve of C57BL/6 mice transplanted with CX-5461+EV-resistant CMB #9 early-passage cells treated with CX-5461 (35 mg/kg thrice weekly) and everolimus (5 mg/kg daily) (n=5-6). Survival curves of drug-naïve cells were actual data reproduced from Devlin et al., 2016. **(C)** Western blot analysis of markers for the on-target effect of 20 nM everolimus treatment for 3 hours, specifically phosphorylation of RPS6 at Serine 240/244 residues and total RPS6 (n=3). Actin was used as a loading control. Synthesis rate of 47S pre-rRNAs in **(D)** drug-naïve (CTRL), **(E)** Everolimus-resistant (EV), **(F)** CX-5461-resistant (CX) and **(G)** CX-5461+EV-resistant (CMB) cell lines in response to 3-hour 50nM CX-5461, or **(H)** CX and **(I)** CMB cell lines in response to 10-hour 50 nM CX-5461 were determined by $^{32}$P-orthophosphate "pulse" labelling and **(J)** its quantitation (n=3). 28S rRNA abundance was used as a loading control. EtBr: ethidium bromide. Data were analysed by Student's t test. Graphs represent mean ± SEM of n=3. *, $P \leq 0.05$. ***, $P \leq 0.001$ **(K)** Western blot analysis for p53 abundance in response to 50 nM CX-5461 treatment for 3 hours. **(L)** Western blot analysis for indicated proteins in response to 5 mM metformin or 20 nM everolimus treatment for 6 hours (n=3). Actin was used as a loading control.

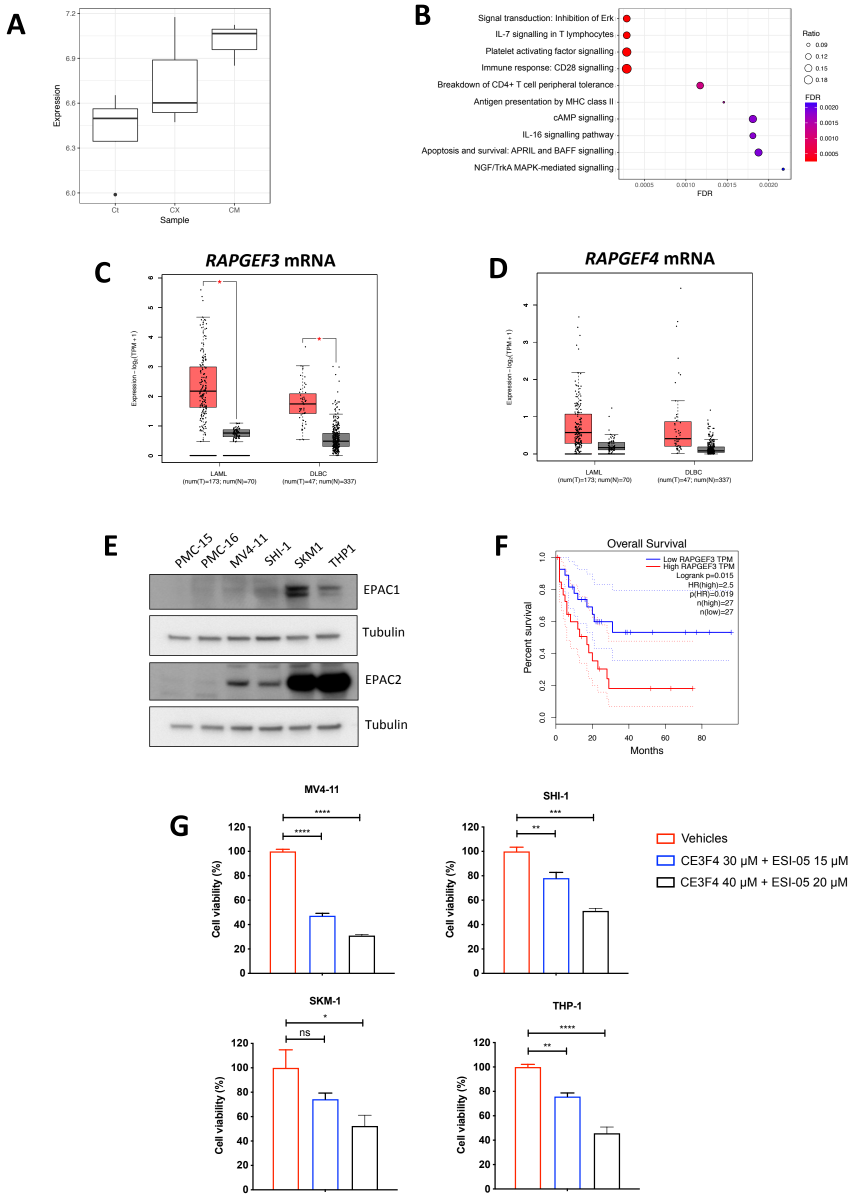


**Fig. S3. Polysome profiling and functional analysis of the early-passage Eμ-Myc lymphoma cell lines**

**(A)** Boxplot showing polysomal abundance of mRNA encoding RAP1 protein in early-passage CTRL, CX and CMB Eμ-Myc B-lymphoma cells. CTRL vs. CX; $p$ = 0.0385. CTRL vs. CMB; $p$ = 0.0081. **(B)** Enrichment analysis by GeneGO MetaCore® of the polysomal RNAseq data comparing CX-5461-resistant (CX) and drug-naïve cells (CTRL) (n=3; false discovery rate (FDR) ≤ 0.05; fold change (FC) ≥ 1.5 or ≤-1.5). Median mRNA expression levels of **(C)** *RAPGEF3* (mRNA encoding EPAC1 protein) and **(D)** *RAPGEF4* (mRNA encoding EPAC2 protein) generated by gene expression profiling interactive analysis (GEPIA) bioinformatics web tool based on The Cancer Genome Atlas (TCGA) database. Red: tumor, black: normal. *, P < 0.05. **(E)** Western analysis of EPAC1 and EPAC2 abundance in white blood cell samples (isolated from healthy donors #PMC-15 and #PMC-16) versus MV4-11, SHI-1, SKM1 and THP1 human AML cell lines (representative of n=2). **(F)** Survival curves of AML patients with low and high EPAC1 (RAPGEF3) (n=27 each; cut-off: 50%) generated by GEPIA based on TCGA database. **(F)** Propidium iodide and **(G)** CellTiterGLO®-based viability assay of MV4-11, SHI-1, SKM-1 and THP-1 cells treated as indicated. Graphs represent mean ± SEM of n=3. Data were analysed by one-way ANOVA. ns, not significant; P≥ 0.05; *, P ≤ 0.05; **, P ≤ 0.01; ***, P ≤ 0.001; ****, P ≤ 0.0001.

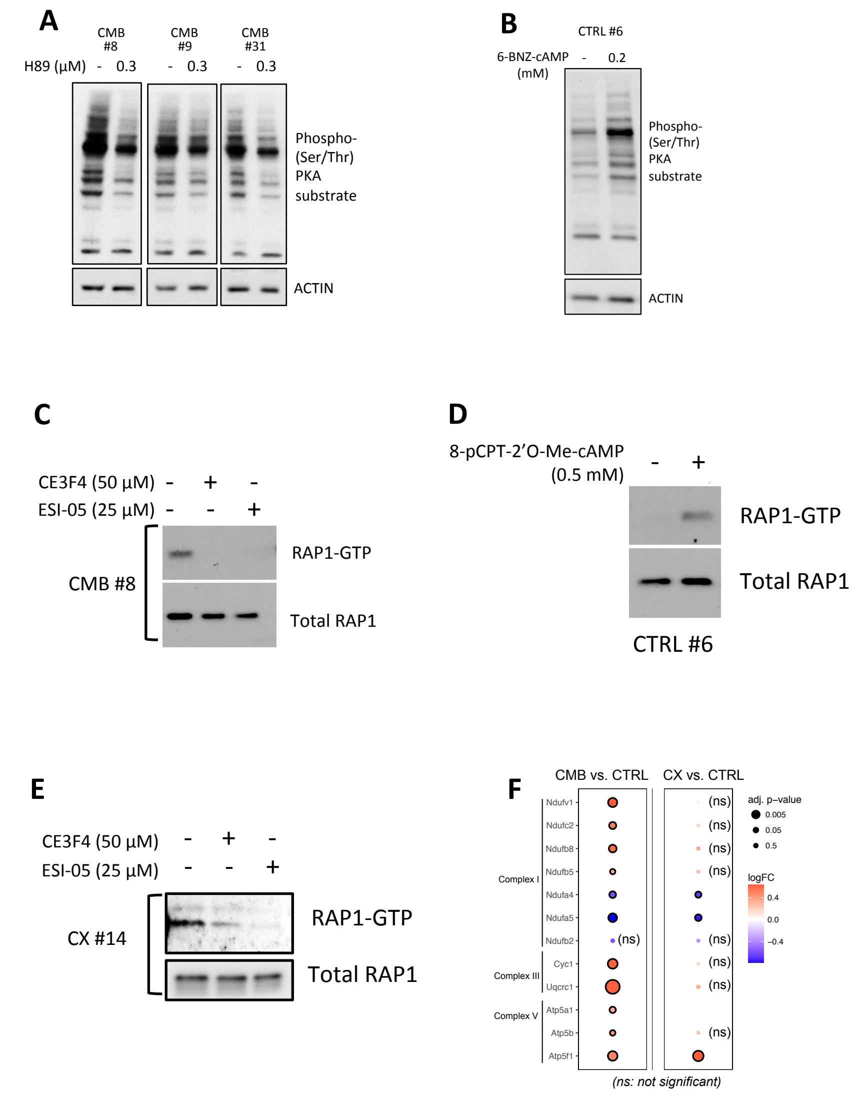


**Fig. S4. Validation of cAMP pathway modulators and expression analysis of components of mitochondrial electron transport chain.**

Western analysis of PKA activity using the phosphorylated PKA substrate antibody on whole cell extracts isolated from the indicated early-passage cell lines treated for 24 hours with **(A)** H89 (n=3) or **(B)** 6-BNZ-cAMP (representative of n=3). Actin was used as a loading control. **(C)** Western analysis of the activated, GTP-bound RAP1 pull-down assays of the early-passage CX-5461+EV-resistant (CMB) cells extracts isolated following treatments with indicated EPAC inhibitors for 24 hours (n=1). **(D)** Western analysis of activated, GTP-bound RAP1 pull-down assays using early-passage drug-naive (CTRL) Eµ-Myc B- cell lymphoma extracts isolated following treatments with the selective EPAC activator 8-pCPT-2-O-Me-cAMP for 24 hours (n=1). **(E)** Western analysis of the activated, GTP-bound RAP1 pull-down assays of the early-passage CX-5461-resistant (CX) cells extracts isolated following treatments with indicated EPAC inhibitors for 24 hours (n=1). **(F)** Polysomal abundance of mRNAs encoding components of mitochondrial oxidative phosphorylation based on polysome profiling data. Red denotes upregulation, blue denotes down regulation (n=3; adjusted P value ≤ 0.05; fold change (FC) ≥ 1.5 or ≤-1.5).

### Supplementary Table

| Protein | Mass (kDa) | Primary Antibody Details | Primary Antibody Dilution | Secondary Antibody Details |
|---|---|---|---|---|
| **Immunoblotting** | | | | |
| β-ACTIN | 42 kDa | MP Biosciences 691002 | 1:30,000 | Goat-α-rabbit HRP[1] |
| p53 | 53 kDa | Leica Biosystems | 1:1,000 | Goat-α-mouse HRP[2] |
| RPS6 | 32 kDa | Cell Signalling #2317 | 1:2,000 | Goat-α-mouse HRP[2] |
| Phospho-RPS6 | 32 kDa | Cell Signalling #2215 | 1:2,000 | Goat-α-rabbit HRP[1] |
| Phospho-AMPKα (Thr172) | 62 kDa | Cell Signalling #2531 | 1:1,000 | Goat-α-rabbit HRP[1] |
| AMPKα | 62 kDa | Cell Signalling #2532 | 1:1,000 | Goat-α-rabbit HRP[1] |
| EPAC1 | 100 kDa | Cell Signalling #4155 | 1:1,000 | Goat-α-mouse HRP[2] |
| EPAC2 | 110 kDa | Abcam #193665 | 1:2,000 | Goat-α-rabbit HRP[1] |
| RAP1 | 21 kDa | Thermo Fisher Scientific #16120 | 1:1,000 | Goat-α-rabbit HRP[1] |
| Phospho-PKA substrate (RRXS*/T*) | N.A | Cell Signalling #9624 | 1:1,000 | Goat-α-rabbit HRP[1] |
| Notes:<br>1. Goat-α-rabbit horseradish peroxide (HRP) IgG (H+L) (Bio-Rad #170-6515); dilution 1:2,000<br>2. Goat-α-mouse horseradish peroxide (HRP) IgG (H+L) (Bio-Rad #170-1011); dilution 1:2,000 | | | | |
| **Flow cytometry** | | | | |
| B220 (CD45R)-APC | N.A. | Thermo Fisher Scientific 17-0452-82 | 1:400 | N.A. |

**Table S1. List of antibodies used in immunoblotting and flow cytometry experiments**
